# A vascular origin for pulmonary smooth muscle in the avian lung

**DOI:** 10.1101/2022.07.13.499952

**Authors:** Aaron H. Griffing, Katharine Goodwin, Michael A. Palmer, Chan Jin Park, Megan Rothstein, Benjamin J. Brack, Jorge A. Moreno, Bezia Lemma, Wei Wang, Ricardo Mallarino, Celeste M. Nelson

## Abstract

Lungs exhibit strikingly diverse epithelial architectures – from the branched airways of mammals to the sac-like lungs of lizards and the looped airways of birds. Across lineages, the pulmonary mesenchyme gives rise to smooth muscle that interacts with and shapes the underlying pulmonary epithelium. In mammals and lizards, pulmonary smooth muscle forms early and drives epithelial branching, whereas in birds it appears only after morphogenesis is largely complete. The developmental basis for this delay has remained unclear. Using comparative single-cell RNA sequencing, ATAC-sequencing, and imaging of mouse, anole, and chicken embryos, we found that smooth muscle in the chicken lung is transcriptionally similar to vascular, rather than visceral, smooth muscle. Strikingly, imaging revealed smooth muscle cells extending between the pulmonary vasculature and the epithelium, and surgical removal of these vessels prevented the formation of smooth muscle around the airways. The vascular transcription factor *PITX2* was highly expressed in these cells and its knockdown markedly reduced smooth muscle differentiation. Taken together, these findings identify vascular smooth muscle as the developmental source of pulmonary smooth muscle in birds and establish *PITX2* as a key regulator of this lineage transition, revealing an unexpected developmental and evolutionary link between the circulatory and respiratory systems.

## INTRODUCTION

The vertebrate lung has diversified extensively to meet the physiological demands of distinct lineages.^1^ In mammals, a hierarchically branched tree-like epithelium terminates in the alveolar gas-exchange surface.^2,3^ Squamates (lizards and snakes) possess a comparatively simple hollow chamber lined by a corrugated epithelial sheet that forms the faveolar gas-exchange surface.^4–6^ Birds, in contrast, evolved a continuous, closed circuit of epithelial airways and auxiliary air sacs that act as bellows,^7,8^ enabling unidirectional flow of air across the gas-exchange surface within air capillaries.^3,7^ Despite these major differences in architecture, lung morphogenesis in all amniotes depends on reciprocal signaling between the pulmonary epithelium and the surrounding mesenchyme.^9–11^ In mammals and squamates, the relatively soft mesenchyme differentiates early into the stiffer pulmonary smooth muscle, which shapes the underlying epithelium into branches or faveolae.^11–16^ By contrast, the avian lung epithelium forms branches in the absence of smooth muscle,^17–19^ which only appears prior to epithelial fusion of the looped airways.^19,20^ The developmental and molecular basis for this delayed onset of smooth-muscle differentiation remains unknown.

Smooth muscle is a contractile tissue present in all organ systems. Visceral smooth muscle lines epithelia such as the gut and lung, whereas vascular smooth muscle forms the walls of blood vessels.^21,22^ In the developing lung, sonic hedgehog (SHH) from the epithelium activates a cascade of transcription factors in the mesenchyme, including myocardin (MYOCD), to promote visceral smooth muscle differentiation by inducing the expression of smooth muscle-specific proteins, including alpha-smooth muscle actin (αSMA).^23–25^ During embryogenesis, distinct progenitor populations – including neural crest and splanchnic mesoderm – also activate MYOCD and transcription factors such as PITX2 to generate vascular smooth muscle.^22,26,27^ Although smooth-muscle differentiation is tightly regulated, it is not terminal: cells retain striking plasticity, switching between contractile and proliferative phenotypes in both embryonic airways^14^ and adult blood vessels.^28^ Vascular smooth muscle cells can even adopt alternative fates, migrating toward the vessel lumen to form foam cells^29,30^ or exiting the vessel wall to contribute to skeletal muscle.^31^ This inherent plasticity raises the possibility that smooth muscle populations might be repurposed during development or evolution. Such flexibility could help explain the unusual timing and origin of smooth muscle in the avian lung, where it appears only after epithelial morphogenesis is largely complete.

To explore this divergence, we compared lung development across three amniote lineages – mouse, anole lizard, and chicken – using immunofluorescence, single-cell RNA-sequencing (scRNA-seq), and assay for transposase-accessible chromatin sequencing (ATAC-seq). These analyses revealed that, unlike mouse and anole lungs where pulmonary smooth muscle differentiates early from the mesenchyme, the chicken lung recruits smooth muscle from a vascular source. Functional perturbations in chicken, including vascular ablation and PITX2 knockdown, confirmed that the developing vasculature is required for pulmonary smooth muscle differentiation. Together, these findings reveal an unexpected developmental link between the vascular and pulmonary smooth muscle in birds.

## RESULTS

To identify when and where pulmonary smooth muscle first emerges across amniotes, we examined smooth muscle differentiation in mouse, anole, and chicken embryos using immunofluorescence for αSMA (**Fig. 1A-B**). In mouse, pulmonary smooth muscle differentiates in a proximal-to-distal wave beginning at embryonic day (*E*) 12.5 (**Fig. 1C**). This wave of differentiation specifies sites of epithelial branching by generating patterns of stiffness around the growing airways.^14,32^ In the brown anole, smooth muscle surrounds the basal surface of the epithelium and subsequently reorganizes into a mesh between 5 and 6 days post-oviposition (dpo) (**Fig. 1D**). The epithelium then deforms though openings in this mesh to generate the characteristic faveolar architecture.^11,16^ In contrast, the embryonic chicken lacks pulmonary smooth muscle during early branching (Hamburger-Hamilton^33^ stages HH23-26), which proceeds through an epithelial-intrinsic mechanism of apical constriction, and only appears at later stages when it wraps extensively around the pulmonary epithelium (HH34, HH38; **Fig. 1E**). Notably, vascular smooth muscle is already present around the pulmonary arteries by HH26. These patterns suggest that, in birds, the timing and source of smooth muscle differentiation differ fundamentally from those of mammals and squamates.

**Figure 1.**
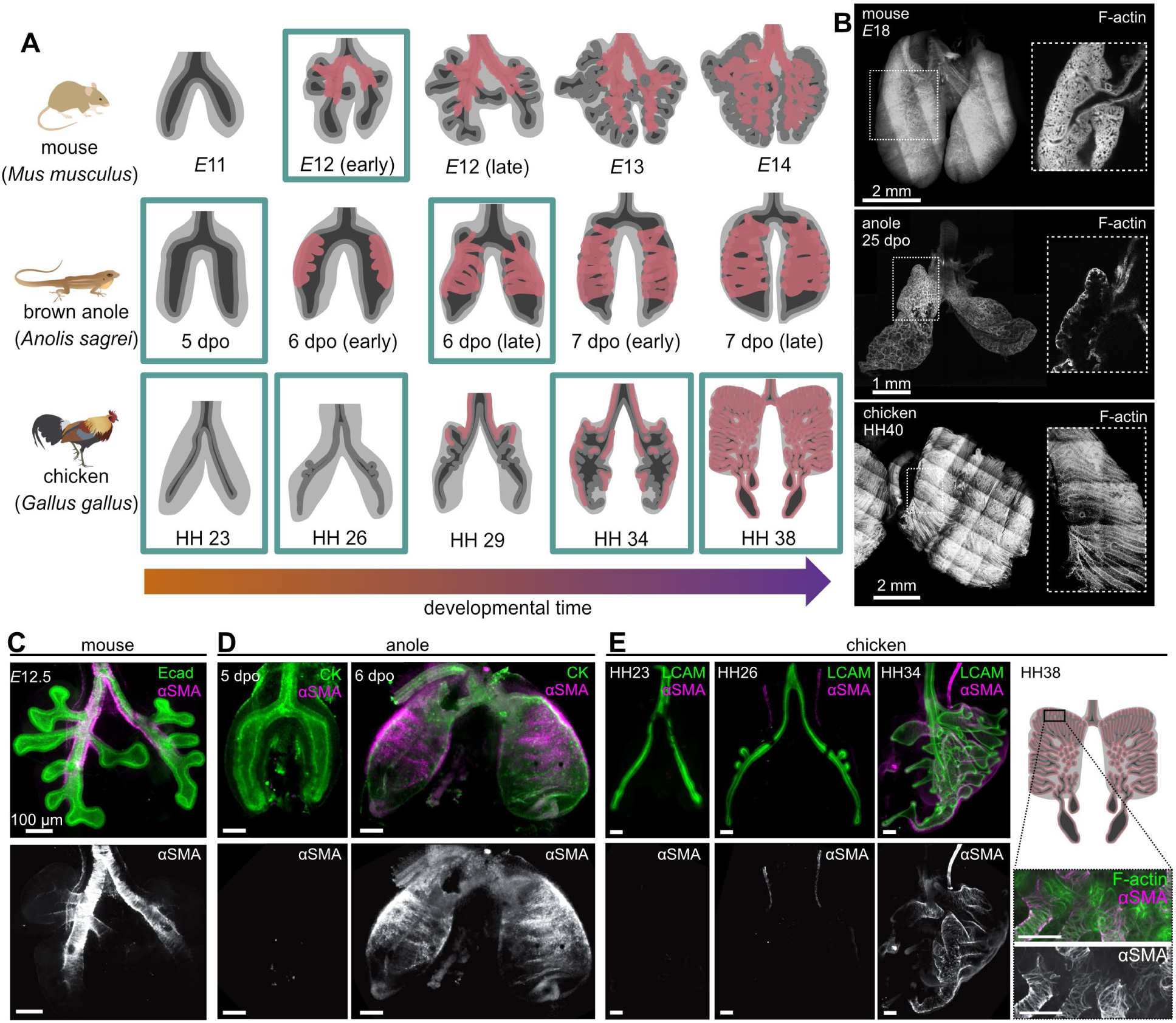
Amniotes exhibit diverse lung morphologies and distinct patterns of smooth muscle differentiation. **A**) Schematic illustrating different stages of lung development in mouse, brown anole, and chicken. Dark grey corresponds to epithelium, light grey corresponds to pulmonary mesenchyme, and pink corresponds to pulmonary smooth muscle. Green boxes denote developmental stages sampled in the present study. **B**) Images show fluorescent staining of F-actin in late embryonic stage mouse, brown anole, and chicken lungs. Scale bars = 1 mm. **C–E**) Images show immunofluorescence for alpha smooth muscle actin (αSMA, magenta) and either E-cadherin (Ecad/LCAM, green, **C, E**) or epithelial cytokeratins (CK, green, **D**). Scale bars = 100 µm.

To characterize these differences molecularly, we profiled lung development across mouse, anole, and chicken embryos at stages encompassing smooth muscle differentiation (**Fig. 2A-C**). Single-cell RNA-seq (scRNA-seq) analysis revealed robust expression of *ACTA2* (αSMA) and *MYOCD* (myocardin) in mesenchymal subsets of mouse (*E*12.5) and anole (6 dpo) lungs, consistent with active formation of pulmonary smooth muscle (**Fig. 2D; Supplemental Figs. 1, 2**). In chicken, however, expression of both smooth muscle markers was sparse across all timepoints (**Fig. 2D; Supplemental Fig. 3**), despite the presence of smooth muscle histologically at HH34–38 (**Fig. 1E**). Bulk ATAC-seq performed on stage-matched embryos revealed open chromatin at the *ACTA2* and *MYOCD* loci in all three species (**Fig. 2E, F**), indicating that the regulatory landscape in the chicken mesenchyme is transcriptionally competent, even though expression of canonical smooth muscle genes remains low. Thus, chromatin accessibility and transcriptional output of smooth-muscle markers are uncoupled in the avian lung.

**Figure 2.**
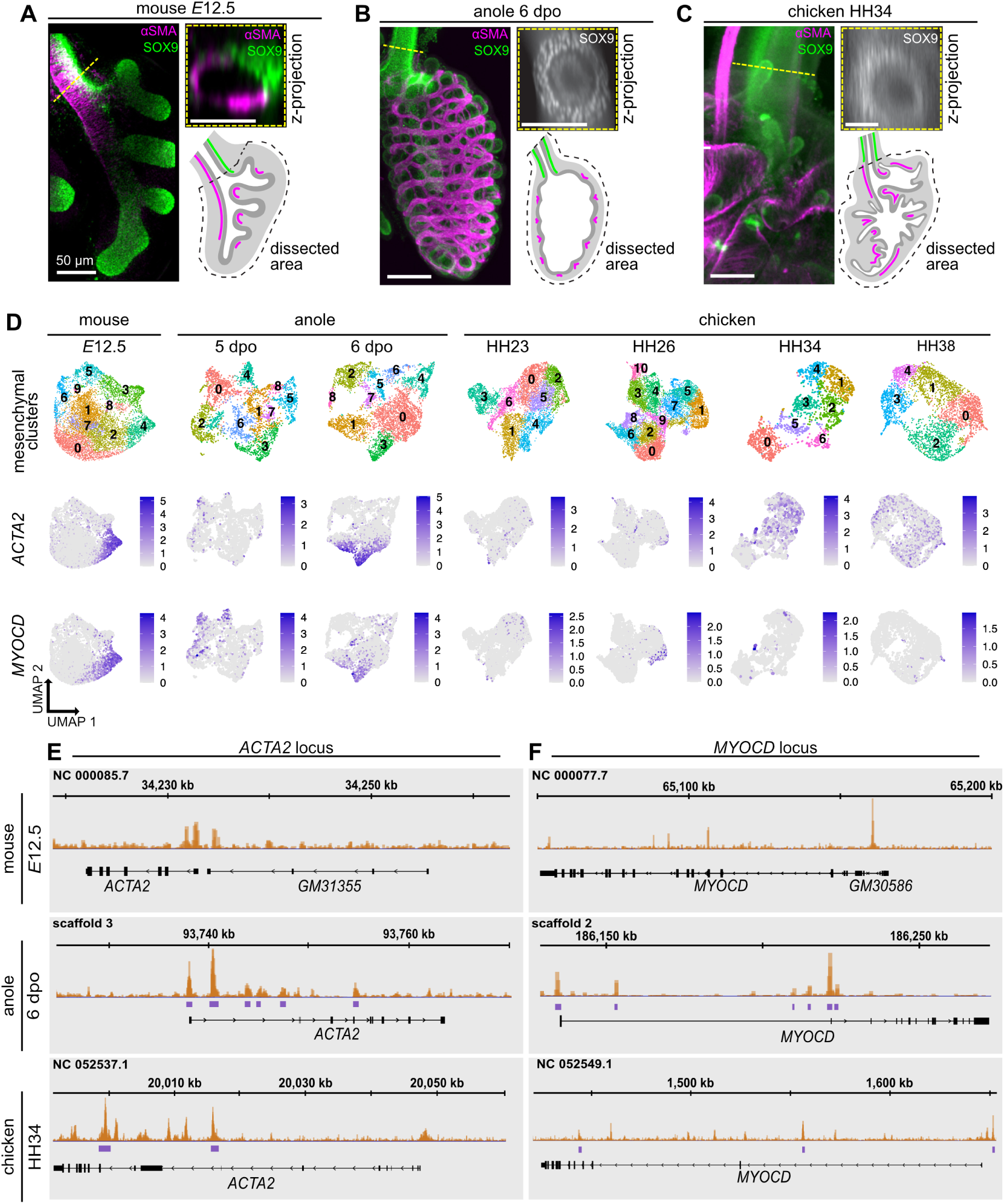
Comparative transcriptomics and genomics of developing amniote lungs illustrate a distinct expression pattern of smooth-muscle markers in chicken. **A–C**) Images show immunofluorescence for αSMA (magenta) and SOX9 (green). Dissected areas correspond to tissue collected for scRNA-seq. Scale bars = 50 µm. **D**) scRNA-seq of developing mouse, anole, and chicken lungs. UMAPs illustrate cells within mesenchymal clusters that express either *ACTA2* or *MYOCD*. **E**) Bulk ATAC-seq of developing mouse, anole, and chicken lungs illustrating open chromatin pileups (orange) and narrow peaks (purple) surrounding regulatory regions of either *ACTA2* or *MYOCD*.

In mammals and squamates, pulmonary smooth muscle differentiates in response to SHH signaling from the embryonic lung epithelium.^11,34^ We therefore asked whether the absence of smooth-muscle markers in the chicken lung reflects attenuated SHH activity. To answer this question, we examined the expression of SHH, its receptor Patched1 (PTCH1), the signal transducer smoothened (SMO), and antagonist hedgehog interacting protein (HHIP). Surprisingly, scRNA-seq analysis revealed that the chicken pulmonary mesenchyme contains cells expressing *PTCH1*, *SMO*, and *HHIP* (**Fig. 3A**, **Supplemental Fig. 4**), comparable to mouse and anole (**Supplemental Figs. 5–6**). Consistently, fluorescence *in situ* hybridization (RNAscope) showed that all three species express *PITCH1* and *HHIP* in the mesenchyme and *SHH* in the adjacent epithelium (**Fig. 3B-D**). These results indicate that the chicken pulmonary mesenchyme possesses the molecular machinery to respond to SHH and differentiate into smooth muscle. To test whether activation of the SHH pathway is sufficient to induce smooth muscle, we treated embryonic chicken lung explants with smoothened agonist (SAG) and assayed for αSMA expression. SAG treatment did not induce αSMA around the airways, which remained devoid of smooth muscle (**Fig. 3E**). Instead αSMA increased in a distinct patch of the distolateral mesothelium and adjacent submesothelial mesenchyme (**Fig. 3E, insets**), a feature also present – though less pronounced – in vehicle-treated control lungs (**Fig. 3E-H**).

**Figure 3.**
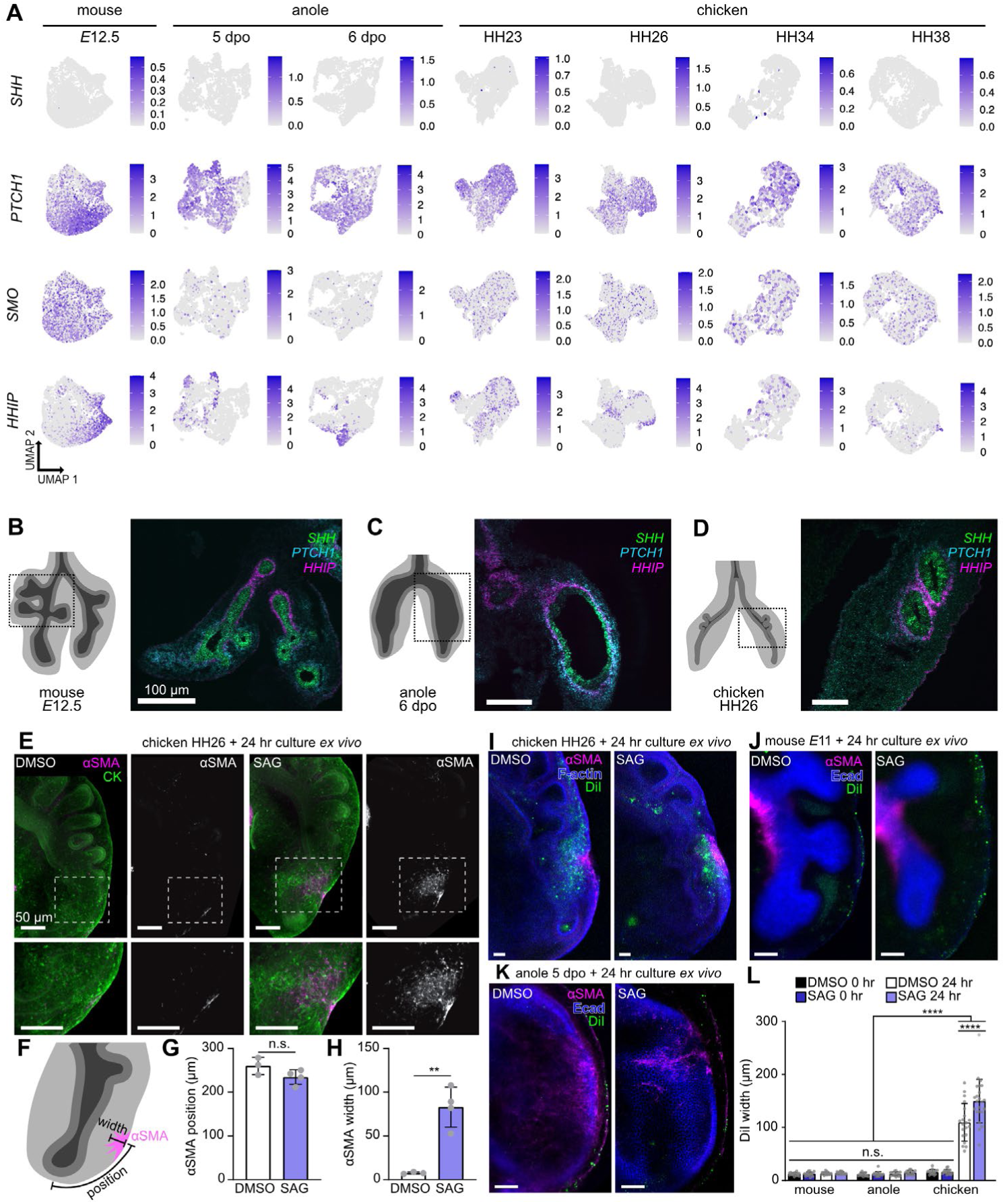
Sonic hedgehog signaling does not induce pulmonary smooth muscle differentiation in the chicken lung. **A**) scRNA-seq of developing mouse, anole, and chicken lungs. UMAPs illustrate cells within mesenchymal clusters that express the sonic hedgehog-pathway genes *SHH*, *PTCH1*, *SMO*, or *HHIP*. **B–D**) Fluorescence *in situ* hybridization of *SHH*, *PTCH1*, and *HHIP* in embryonic lungs of mouse (**B**), anole (**C**), and chicken (**D**). Scale bars = 100 µm. **E**) Images show immunofluorescence for αSMA (magenta) and epithelial cytokeratins (CK, green) in HH26-stage chicken lungs treated with either DMSO or SAG for 24 hours. Scale bars = 50 µm. **F**) Schematic illustrating how width and position of αSMA+ cells were measured. **G–H**) Quantification of width and position of the cluster of αSMA+ cells in HH26-stage chicken lungs treated with DMSO or SAG for 24 hours. **I–K**) Images show immunofluorescence for αSMA (magenta), DiI labeling (green), and either staining for F-actin (blue, **I**) or immunofluorescence for E-cadherin (Ecad, blue, **J**, **K**) in HH26-stage chicken lungs (**I**), *E*11 mouse lungs (**J**), or 5-dpo anole lungs (**K**) treated with either DMSO or SAG for 24 hours. Scale bars = 50 µm. **L**) Quantification of the width of the cluster of DiI-labeled cells before and after culture in the presence of DMSO or SAG in all three species.

These findings suggested that a subset of mesothelial cells responds to SHH by migrating into the underlying pulmonary mesenchyme. To test this hypothesis directly, we labeled the mesothelium with DiI and cultured embryonic lung explants with or without SAG. After 24 hours of culture, DiI-labeled cells were present within the mesenchyme adjacent to the distolateral mesothelium in chicken lungs (**Fig. 3I**, **Supplemental Fig. 7**), but remained restricted to the mesothelium in mouse and anoles (**Fig. 3J-L**, **Supplemental Fig. 7**). Thus, despite conserved expression of SHH-pathway components, the chicken pulmonary mesenchyme exhibits a distinct response, redirecting SHH-induced differentiation to a mesothelial-derived population.

Given the distinct response of the chicken lung to SHH signaling, we hypothesized that its pulmonary smooth muscle might represent a fundamentally different cell type from that of mouse or anole. To test this hypothesis, we compared the transcriptomes of *ACTA2*+/*MYOCD*+ cells from each species with those of different contractile lineages from mouse, including pulmonary smooth muscle, myofibroblasts, pericytes, and vascular smooth muscle from embryonic and postnatal lungs,^35^ myocardial cells from embryonic hearts,^36^ lymphatic smooth muscle from adult vessels,^37^ and skeletal muscle stem cells from adult quadriceps and diaphragm^38^ (**Fig. 4A-C**). As expected, the *ACTA2*+/*MYOCD*+ cluster from mouse (*E*12.5, cluster 4) most closely matched pulmonary smooth muscle cells from the reference dataset (**Fig. 4A**). The corresponding cluster from anole (6 dpo, cluster 3) also aligned with pulmonary smooth muscle from mouse (**Fig. 4B**). In contrast, the chicken *ACTA2*+/*MYOCD*+ clusters (HH34, clusters 1 and 5) showed strongest similarity to vascular smooth muscle from *E*15 mouse lungs (**Fig. 4C**). These cross-species comparisons reveal that pulmonary smooth muscle in the chicken is transcriptionally aligned with vascular smooth muscle, suggesting a fundamental shift in tissue identity relative to mammals and squamates.

**Figure 4.**
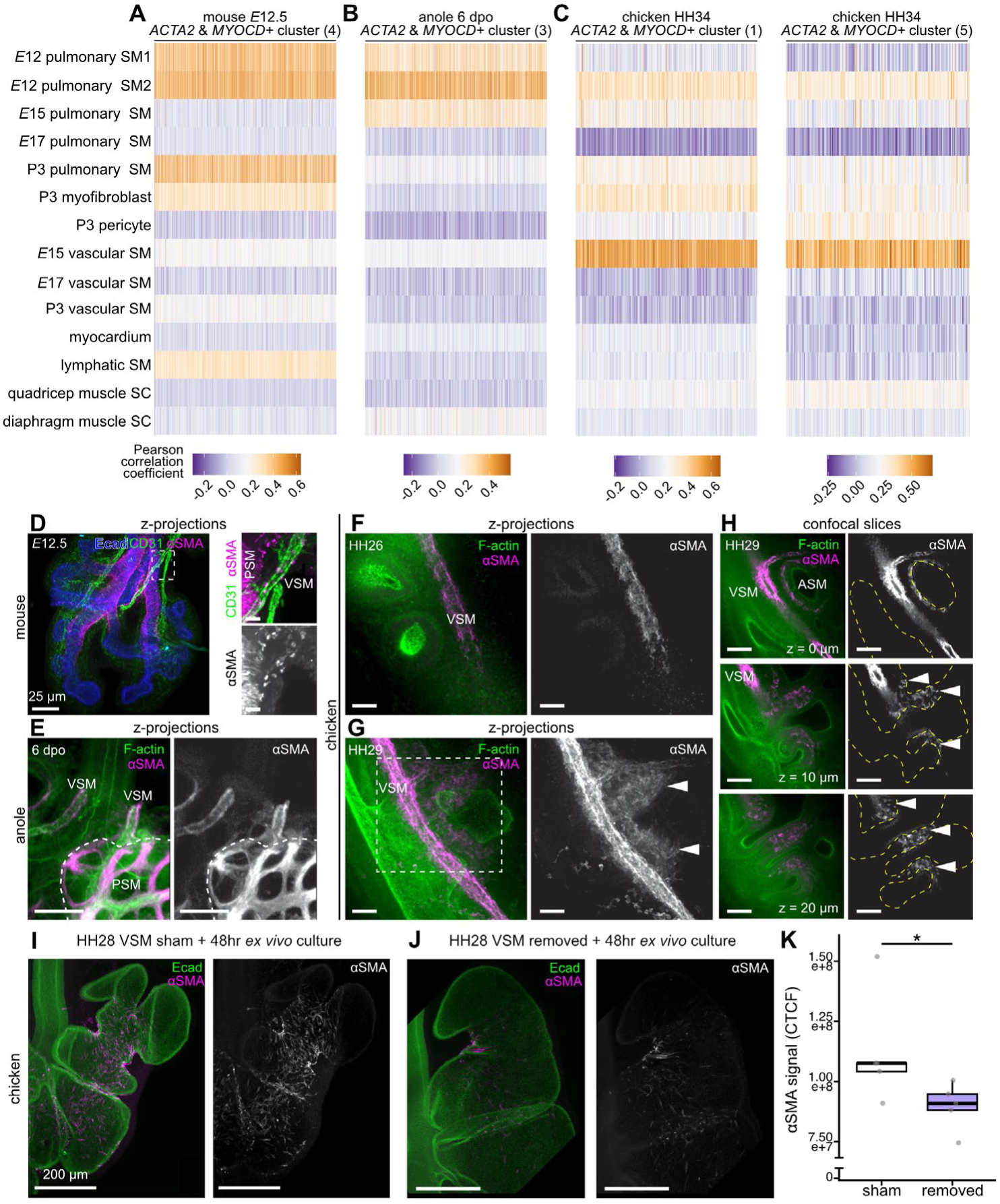
Pulmonary smooth muscle in chicken lungs appears to be derived from vascular smooth muscle. **A–C**) Pearson correlation analysis comparing transcriptional similarity between *ACTA2*+ *MYOCD*+ clusters from *E*12.5 mouse (**A**), 6-dpo anole (**B**), or HH34-stage chicken (**C**) and previously published mouse datasets. **D**) Images show immunofluorescence for αSMA (magenta), CD31 (green), and Ecad (blue) in an *E*12.5 mouse lung. Zoomed-in images correspond to region in dashed white box. Pulmonary smooth muscle (PSM) and vascular smooth muscle (VSM) are indicated. **E**) Images show immunofluorescence for αSMA (magenta) and F-actin (green) in a 6-dpo anole lung. PSM and VSM are indicated. **F–H**) Images show immunofluorescence for αSMA (magenta) and F-actin (green) in HH26-(**F**) and HH29-stage chicken lungs (**G–H**). Yellow dashed lines outline epithelium. White arrowheads indicate αSMA+ cells that appear to be migrating through the mesenchyme between the blood vessel and airway epithelium. Scale bars = 25 mm. **I–J**) Images show immunofluorescence for αSMA (magenta) and Ecad (green) in the first branch region of vascular smooth muscle sham (**I**) and removal (**J**) lungs after *ex vivo* culture for 48 hours. **K**) Quantification of αSMA signal (corrected total cellular fluorescence, CTCF) in vascular smooth muscle sham and removal experiments following 48 hours of culture *ex vivo*.

The transcriptional similarity between chicken pulmonary smooth muscle and vascular smooth muscle prompted us to examine vascular smooth muscle organization across species using immunofluorescence for αSMA and the endothelial marker CD31. At the stages profiled by scRNA-seq and ATAC-seq, mouse and anole lungs contained only limited vascular smooth muscle restricted to large blood vessels running parallel to the upper airways (**Fig. 4D-E**). In both species, the amount of pulmonary smooth muscle far exceeded that of vascular smooth muscle at these stages of development. In contrast, smooth muscle in HH26-stage chicken lungs was exclusively vascular, encircling the pulmonary arteries adjacent to the airways (**Fig. 4F**). By HH29, αSMA+ cells extended radially outward from this vascular smooth muscle layer into the surrounding mesenchyme (**Fig. 4G**). These αSMA+ cells subsequently filled the interstitial mesenchymal space between blood vessels and epithelial branches (**Fig. 4G**), and small clusters began to wrap the epithelium near the pulmonary artery (**Fig. 4H**). We also observed these spatiotemporal patterns of αSMA+ cells in lungs of the duck (**Supplemental Fig. 8**). Taken together, these observations support the intriguing possibility that, in birds, pulmonary smooth muscle originates from vascular smooth muscle that migrates into the pulmonary mesenchyme.

We next tested whether vascular smooth muscle is required for the formation of pulmonary smooth muscle in the embryonic chicken lung. We isolated lungs at HH28 and cultured them *ex vivo* after surgically removing the major vessels running parallel to the primary bronchi. In sham controls with intact vasculature, αSMA+ cells surrounded the airway epithelium by the end of the culture period (**Fig. 4I**). In contrast, explants lacking vasculature displayed a marked reduction of αSMA+ cells around the airway epithelium (**Fig. 4J-K**). These results demonstrate that vascular smooth muscle is necessary for the establishment of pulmonary smooth muscle in the developing chicken lung.

To identify molecular regulators of this vascular-to-pulmonary transition, we leveraged our chicken scRNA-seq dataset by integrating mesenchymal clusters across all four developmental stages. This analysis revealed two *ACTA2+* populations, only one of which also expressed *MYOCD* (**Fig. 5A**). The top marker for this *ACTA2*+/*MYOCD*+ cluster was *PITX2* (**Fig. 5A**), a homeodomain transcription factor previously found to regulate the early stages of vascular smooth muscle differentiation and vascular patterning.^26,39,40^ Expression of *PITX2* was minimal in mouse and anole (**Fig. 5B**), consistent with the limited amounts of vascular smooth muscle at the stages analyzed (**Fig. 4D, E**). In contrast, *PITX2* expression was strongly enriched in the chicken (**Fig. 5B**), particularly during earlier stages exhibiting robust development of vasculature (**Fig. 4F**). Consistently, ATAC-seq further showed open chromatin at the *PITX2* locus in early chicken lungs (**Supplemental Fig. 9**), supporting transcriptional activation of this vascular program.

**Figure 5.**
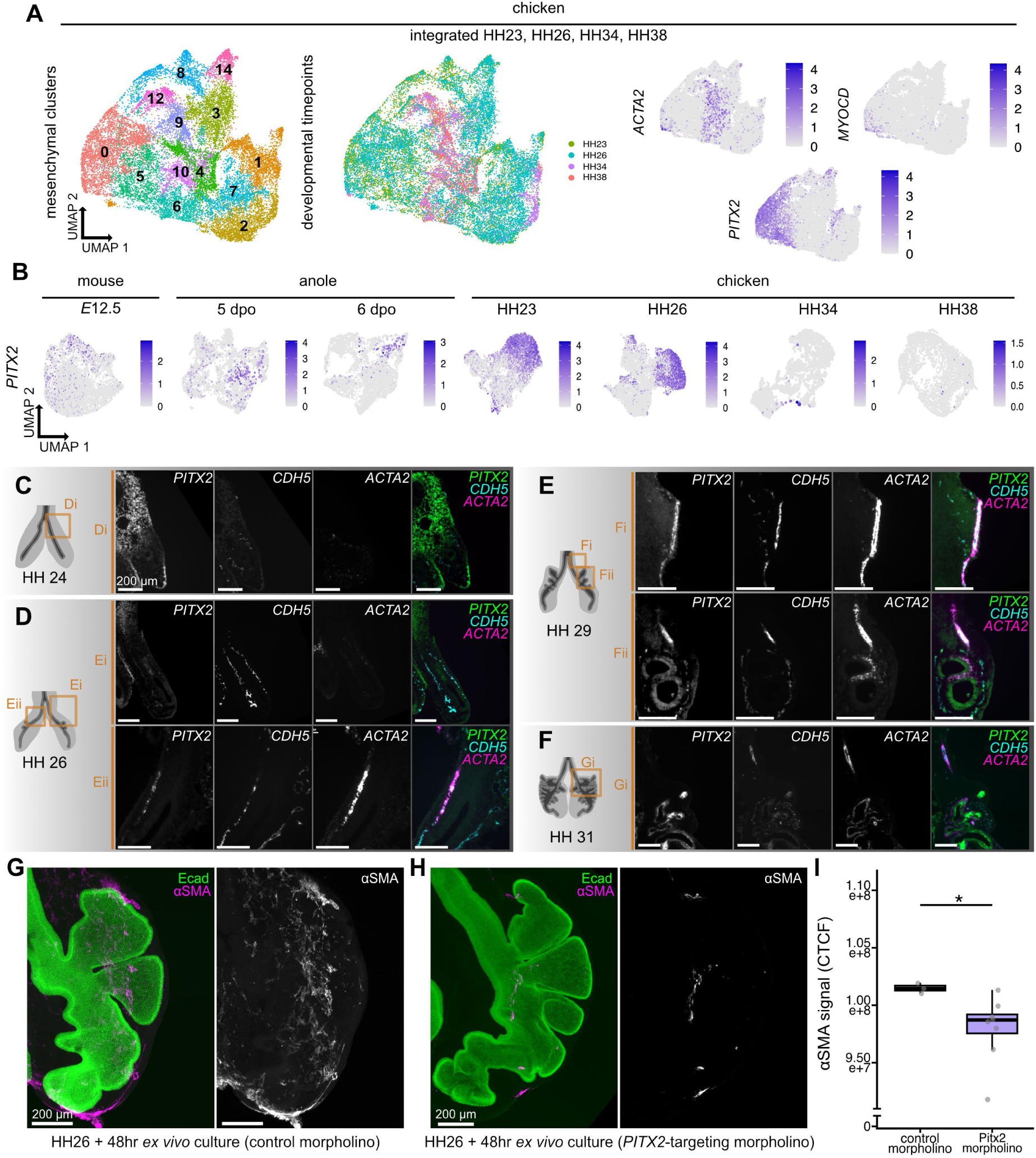
*Pitx2* expression and regulation in the pulmonary mesenchyme. **A**) UMAPs illustrating an integrated dataset of HH23-, HH26-, HH34-, and HH38-stage mesenchymal cells and expression of either *ACTA2*, *MYOCD*, or *PITX2*. **B**) UMAPs illustrating *PITX2* expression in developing mouse, anole, and chicken lung mesenchyme. **C**–**F**) Fluorescence *in situ* hybridization of *PITX2*, *CDH5*, and *ACTA2* in embryonic chicken lungs at HH24 (**C**), HH26 (**D**), HH29 (**E**), and HH31 (**F**). Scale bars = 200 µm. **G–I**) Images show immunofluorescence for αSMA (magenta) and Ecad (green) in lungs exposed to either control morpholino (**G**) or *PITX2*-targeting morpholino (**H**) and then cultured *ex vivo* for 48 hours. Scale bars = 200 µm. **I**) Quantification of αSMA signal (corrected total cellular fluorescence, CTCF) in morpholino experiments following 48 hours of culture *ex vivo*.

To confirm the identity of the *PITX2*-expressing population in the embryonic chicken lung mesenchyme, we conducted RNAscope for *PITX2*, *ACTA2*, and the vascular endothelial marker, *CDH5* (which encodes for VE-cadherin). At early stages (HH24 and HH26), *PITX2* was expressed in the proximal pulmonary mesenchyme (**Fig. 5C, D**) and in *ACTA2*+ cells adjacent to *CDH5*+ endothelium (**Fig. 5D**). By HH29, *PITX2* remained expressed in *ACTA2*+ cells adjacent to endothelium but also appeared in those surrounding the airway epithelium (**Fig. 5E**). At this stage, *PITX2* was no longer expressed in the proximal mesenchyme, consistent with our scRNA-seq data (**Fig. 5B**), and began to localize within the epithelium itself. By HH31, *PITX2* expression was largely confined to the epithelium (**Fig. 5F**). Together, these data indicate that *ACTA2*+/*PITX2*+ cells initially correspond to vascular smooth muscle, a subset of which migrates and becomes pulmonary smooth muscle around the airways.

To test whether *PITX2* is required for this process, we downregulated its expression in HH28-stage lung explants by injecting targeted morpholinos into the proximal pulmonary mesenchyme (**Supplemental Fig. 10**). Explants injected with control morpholinos developed extensive αSMA+ smooth muscle throughout the lung after 48 hours in culture (**Fig. 5G**). In contrast, explants injected with *PITX2*-targeting morpholinos showed a significant attenuation of αSMA+ cells (**Fig. 5H-I**). These results demonstrate that PITX2 is required for smooth muscle differentiation and, consequently, for the formation of pulmonary smooth muscle in the embryonic chicken lung.

## DISCUSSION

Morphogenesis of the vertebrate lung involves interactions between the pulmonary epithelium and its adjacent smooth muscle.^11,12,20^ Given that the tissues required for lung morphogenesis are conserved across vertebrates, one might expect conservation in the underlying gene-expression patterns and morphogenetic processes. However, our results reveal that the chicken pulmonary mesenchyme exhibits a distinct transcriptomic landscape and responds differently to SHH signaling compared with its mammalian and reptilian counterparts.^11,15^ These interspecific differences are reminiscent of patterns observed in other endoderm-derived organs. For instance, intestinal villus formation proceeds independently of smooth muscle differentiation in the mouse but occurs concurrently with it in the chicken.^41,42^ In this context, SHH signaling promotes smooth muscle differentiation in the mouse yet inhibits it in the chicken.^43^ This recurring contrast suggests that developmental programs governing mesenchymal differentiation and SHH responsiveness have been reconfigured between mammals and birds, potentially across multiple organ systems.

Vascular smooth muscle originates from several different tissues during embryonic development, including the neural crest, proepicardial mesothelium, somitic mesoderm, splanchnic mesoderm, and endothelial cells.^22,44,45^ By contrast, the origins of pulmonary smooth muscle remain poorly characterized, reflecting the relative inattention given to visceral mesenchyme in general, and the pulmonary mesenchyme in particular. In the mouse, pulmonary smooth muscle cells differentiate from progenitors within the sub-epithelial mesenchyme,^14,46,47^ although fate-mapping studies have revealed minor contributions from the distal pulmonary mesenchyme,^48^ neural crest,^49^ and mesothelium.^50^ Together with our findings, these observations suggest that the pulmonary smooth muscle of ancestral vertebrates may have arisen from similarly diverse progenitor populations as those giving rise to vascular smooth muscle. Elucidating the developmental origins of pulmonary (and other visceral) smooth muscle across vertebrate lineages will be key to understanding how these programs have diversified throughout evolution.

Our data uncover a previously unrecognized contribution of vascular smooth muscle, or its progenitors, to the smooth muscle layer that surrounds the avian airway epithelium. This discovery raises several mechanistic possibilities. One is that vascular smooth muscle cells undergo transdifferentiation into pulmonary smooth muscle, reminiscent of the conversion of vascular smooth muscle into macrophage-like cells during mammalian atherogenesis.^51^ Alternatively, pulmonary smooth muscle mat differentiate in situ within a perivascular niche adjacent to pulmonary vessels. Pericytes within such niches in other developing organs express smooth-muscle markers^52^ and can serve as multipotent progenitors capable of generating skeletal muscle.^53^ A third possibility is that pulmonary smooth muscle precursors arise outside the lung and migrate along the blood vessels to reach their destination. Recent advances in replication-incompetent avian retroviruses,^54^ CRISPR-Cas9 delivery systems,^55^ and avian embryonic stem cell derivation^56^ now make it feasible to distinguish between these scenarios and dissect the cellular and molecular mechanisms underlying this process.

## METHODS

### Animals and isolation of embryonic lungs

Lungs were dissected from embryos of all species in sterile phosphate-buffered saline (PBS) using either forceps or fine tungsten needles (Fine Science Tools). Mouse embryos were obtained from a captive colony of CD1 mice (*Mus musculus*) at embryonic day (*E*) 12.5. Anole embryos were obtained from a captive colony of wild-caught brown anoles (*Anolis sagrei*). Anole eggs were collected daily and subsequently incubated in a benchtop incubator at 28°C and 80% humidity until the desired stages of 5 and 6 days postoviposition (dpo). Chicken embryos were obtained from fertilized chicken (*Gallus gallus*) eggs, commercially acquired from either Moyer’s Chicks (Quakertown, PA) or University of Connecticut Poultry Farm (Storrs, CT) and incubated at 38°C in a humidified chamber until the desired Hamburger & Hamilton (HH) stages of HH23, HH26, HH34, and HH38.^33^ Duck embryos were obtained from fertilized Pekin duck (*Anas platyrhynchos*) eggs that were commercially acquired from Metzer Farms (Gonzales, CA) and incubated at 38°C in a humidified chamber until the desired stages of HH26, HH27, and HH28. Breeding of mice and anoles and isolation of embryos was carried out in accordance with institutional guidelines following the NIH Guide for the Care and Use of Laboratory Animals and approved by Princeton University’s Institutional Animal Care and Use Committee (IACUC protocols: 2152F, 2104, 1934A, 1855).

### Genetic nomenclature

There are different standards of genetic nomenclature between mouse,^57^ anole,^58^ and chicken. To avoid confusion between these naming conventions, we used the conventions of the Chicken Gene Nomenclature Consortium.^59^

### Single-cell RNA sequencing (scRNA-seq)

Lungs were isolated from embryos and then dissociated through digestion in dispase II (20 mg in 10 mL of DMEM/F12 medium) at room temperature for 10 minutes, with additional mechanical dissociation using fine tungsten needles to generate single-cell suspensions. Dispase was then inactivated by adding DMEM/F12 medium (without HEPES or phenol red; ThermoFisher) supplemented with 5% heat-inactivated fetal bovine serum (FBS; Atlanta Biologicals) and antibiotics (50 units/ml of penicillin and streptomycin). Cell suspensions were next passed through 40-μm-diameter mesh filters. scRNA-seq library preparation and sequencing were then carried out by the Princeton Genomics Core

Facility. Briefly, cells were loaded and processed using the Chromium Single Cell 3’ Library and Gel Bead Kit v3 on the Chromium Controller (10X Genomics) following manufacturer protocols. Individual cells were encapsulated in droplets with individual gel beads carrying unique barcoded primers and lysed. Then, cDNA fragments were synthesized, barcoded, and amplified by PCR. We prepared Illumina sequencing libraries from the amplified cDNA from each sample group using the Nextra DNA library prep kit (Illumina). Libraries were sequenced on NovaSeq 6000 S Prime flowcells (Illumina) as paired-end 28 + 94 nucleotide reads, following manufacturer protocols. We performed base calling and filtered raw sequencing reads using the Illumina sequencer control software to keep only pass-filtered reads for downstream analysis. CellRanger^60^ (v7.1.0) was used to run the count pipeline with default settings on all FASTQ files from each sample to generate gene-barcode matrices using the *M. musculus* reference genome (GRCm39; NCBI PRJNA20689), *A. sagrei* reference genome (AnoSag2.1^61^), or *G. gallus* reference genome (GRCg7b; NCBI PRJNA660757). Finally, CellBender^62^ (v0.3.2) was used to remove background noise from all samples.

scRNA-seq data were exported from CellBender, imported into R^63^ (v4.4.2), and then processed and analyzed using Seurat^64^ (v5). Datasets were normalized and integrated using the Seurat function FindVariableFeatures based on 2000 genes. Datasets were then scaled before analysis using the Seurat functions FindNeighbors and FindClusters to find neighbors and clusters. Finally, clusters were visualized using the uniform manifold approximation and projection (UMAP) dimensional reduction. Initial clustering was carried out on all cells, and then mesenchymal and smooth-muscle-cell clusters were computationally isolated for further analysis. For the integrated analysis, all timepoints of the chicken data were integrated in Seurat using SCTransform^65^ and then processed using Seurat as outlined above.

For Pearson correlation analysis, eight published scRNA-seq datasets were downloaded from GEO, representing embryonic and postnatal mouse lungs (GSE149563^35^), embryonic mouse hearts isolated at *E*7.75, *E*8.25, and *E*9.25 (GSE126128^36^), lymph node stromal cells from adult mice (GSE112903^37^), and quadricep and diaphragm muscle from adult mice (GSE138707^38^). All datasets were processed following the standard Seurat pipeline and relevant clusters were identified based on markers used in the original publications. In the lung datasets, *ACTA2* with *HHIP* were used as markers for pulmonary smooth muscle, *ACTA2* with *NOTCH3* for vascular smooth muscle, *ACTA2* with *STC1* for myofibroblasts, and *ACTA2* with *PDGFRB* for pericytes.^35^ In the heart dataset, *ACTA2*, *MYL9*, and *TTN* were used as markers to identify myocardial cells.^36^ In the lymph node stromal cell dataset, *ACTA2*, *MYH11*, and *MYL9* were used as markers to identify lymphatic smooth muscle cells.^37^ In the muscle datasets, which contained only mononucleated cells, *MYOD1* and *PAX7* were used as markers to identify muscle stem cells.^38^ For each dataset, average expression values of variable genes (based on the Seurat pipeline) were obtained for the cluster(s) of interest. These lists were then filtered based on whether genes were also detected in smooth muscle cells of this study, and the top 50 markers per list based on highest expression level were selected. Finally, Pearson correlation analysis was performed using these top 50 markers for each of the *ACTA2*+ and *MYOCD*+ cell clusters from this study (*E*12.5 mouse – cluster 4, 6-dpo anole – cluster 3, and HH34-stage chicken – clusters 1 and 5).

### Assay for transposase-accessible chromatin sequencing (ATAC-seq)

Lungs were isolated from embryos at the stages of interest and three biological replicates were acquired for each species and developmental stage. Due to drastic differences in lung size between species and developmental stages, different numbers of lungs were pooled for each replicate. For *E*12.5 mouse, 5-dpo anole, 6-dpo anole, HH23-stage chicken, and HH26-stage chicken, 5 lungs were pooled per replicate. For HH34-stage chicken, 4 lungs were pooled per replicate. For HH38-stage chicken, 1 lung was used per replicate. Following isolation, each sample was dissociated through digestion in 0.2% dispase in DMEM/F12 medium at 37°C for 60 minutes, with additional mechanical dissociation using fine tungsten needles to generate single-cell suspensions. Dispase solution was inactivated by adding DMEM/F12 medium supplemented with 5% heat-inactivated FBS. The remaining cell suspension was then passed through 40-μm-diameter mesh filters, washed with 0.2% bovine serum albumin (BSA) in PBS, and centrifuged at 500 rcf for 10 minutes at 4°C. Supernatant was discarded and the pellet was resuspended in 0.2% BSA in PBS and finally passed through an additional mesh filter. Cell viability was determined using Trypan Blue (Sigma), after which a solution of 1,000 cells/µL was used for library preparation.

ATAC-seq library preparation was carried out by the Princeton Genomics Core Facility following Omni-ATAC methods of Moreno et al.^66^ ATAC-seq libraries were sequenced on Illumina NovaSeq 6000 S Prime as pair-end 61-nt reads. Raw sequencing reads were filtered by the NovaSeq control software and Pass-Filter reads were used for downstream analyses. Reads were trimmed using NGmerge^67^ and mapped to either the *M. musculus* reference genome (GRCm39; NCBI PRJNA20689), *A. sagrei* reference genome (AnoSag2.1^61^), or *G. gallus* reference genome (GRCg7b; NCBI PRJNA660757) using Bowtie2.^68^ For each biological replicate, PCR duplicates were removed using Picard (Broad Institute) and reads were multi-mapped and filtered using Samtools.^69^ Effective genome size was estimated for each species using faCount^70^ and subsequently peaks were called on each biological replicate using MACS3.^71^ Irreproducible Discovery Rate^72^ (IDR) was used to compare peak calls between replicates and only narrow peaks passing a false discovery threshold of 0.05 were further used.

### Immunofluorescence analysis

Lungs were fixed in 4% paraformaldehyde in PBS for either 15 minutes at room temperature or 30 minutes at 4°C and then permeabilized with 0.1% Triton X-100 in PBS. Prior to staining, lungs were blocked with 0.1% BSA and 5% goat serum in PBS with 0.1% Triton X-100. Lungs were then incubated overnight with primary antibodies against αSMA (Sigma a5228, 1:400 or Abcam ab5694, 1:200), CD31 (Abcam ab28364, 1:200), epithelial cytokeratins (CK; Dako Z0622, 1:400), E-cadherin (Invitrogen 13-1900, 1:200), LCAM (DHSB 7D6, 1:50), PITX2 (Abnova H00005308-M01, 1:200), and/or SOX9 (Abcam ab26414, 1:1000) diluted in blocking buffer, followed by incubation with Alexa Fluor-conjugated secondary antibodies (1:200) or phalloidin (1:500) and Hoechst (1:1000). Lungs were then dehydrated in a methanol series (or isopropanol series for phalloidin-stained samples) and then cleared in Murray’s Clear (1:2 benzyl alcohol:benzyl benzoate) or cleared without dehydration in serial dilutions of glycerol (25, 50, 75, and 100%). Lung sections were mounted in Fluoromount (Thermo Fisher). Lungs were imaged on a Hamamatsu MAICO MEMS confocal unit fitted to an inverted microscope (Nikon Eclipse T*i*) or a Crest Optics X-Light V2tp confocal unit fitted to an inverted microscope (Nikon Eclipse T*i*2) with either 10× air or 20× air objectives.

### Fluorescence *in situ* hybridization

Lungs were fixed in 4% paraformaldehyde in PBS for 24 hours at 4°C, and then washed in PBS, 20% sucrose in PBS, and 30% sucrose in PBS for 1-hour increments at 4°C. Lungs were then embedded in optimal cutting temperature (OCT) medium (Tissue Tek), frozen rapidly in a bath of 100% ethanol and dry ice, and stored at −80°C prior to sectioning. Lungs were coronally sectioned at 12-µm thickness and adhered to Superfrost Plus slides (Fisherbrand). Fluorescence *in situ* hybridization was performed using the RNAScope Multiplex Fluorescent V2 Assay (ACD) protocol for fixed-frozen samples.^73^ Probes were used to target *SHH* (mouse, 314361; anole, 882311; chicken, 551581), *PTCH1* (mouse, 402811-C2; anole, 1154541-C2; chicken, 551571-C2), *HHIP* (mouse, 448441-C3; anole, 1285971-C3; chicken, 1285991-C3), *PITX2* (chicken, 499221), *ACTA2* (chicken, 1169151-C2), and *CDH5* (chicken, 458221-C3). Slides were imaged using a Crest Optics X-Light V2tp confocal unit fitted to an inverted microscope (Nikon Eclipse T*i*2) with either 10× air or 20× air objectives.

### Ex vivo culture

Lungs were isolated from mouse, anole, or chicken embryos at the indicated stages and cultured at the air-liquid interface atop porous filters floating on DMEM/F12 medium (without HEPES) supplemented with 5% heat-inactivated FBS and antibiotics. Explants were incubated for either 24 or 48 hours at 37°C under 5% CO_2_. For DiI experiments, lungs were submerged in PBS with 5% Vybrant DiI cell labeling solution (Invitrogen V22885) for 20 minutes at 37°C. To activate SHH signaling, the culture medium was supplemented with 1-μg/ml smoothened agonist (SAG; Calbiochem).

### Surgical removal and morpholino experiments

For surgical removal experiments, chicken lungs were isolated at HH28 and submerged in PBS. Sham replicates (control; N=5) immediately proceeded to *ex vivo* culture for 48 hours. For experimental replicates (N=5), vasculature was removed via dissection using #5 forceps (Fine Science Tools). Following dissection, experimental replicates were cultured *ex vivo* for 48 hours.

For electroporation experiments, chicken lungs were isolated at HH26 and submerged in Ringer’s solution.^74^ Control (GeneTools LLC, PCO-StandardControl-300-F) and targeted morpholinos (GeneTools LLC, AUG-PITX2-201, AUG-PITX2-202) were delivered to the lung mesenchyme via injection using 1-mm-diameter pulled glass needles. Both control and *PITX2*-targeting morpholinos were injected at a final concentration of 0.5 mM, supplemented with 1 µg/µL of carrier DNA and 10-mM Tris pH 8.0. For targeted morpholinos, a cocktail of two morpholinos was generated to target all possible isoforms of chicken *PITX2*. Immediately following injection, lungs were electroporated by placing them between two platinum electrodes (submerged in Ringer’s solution) and delivering five 50-ms pulses of 18V, with an interval of 100-ms between pulses. Following electroporation, lungs were cultured *ex vivo* for 48 hours.

### Image analysis and statistics

Image analysis was carried out in Fiji.^75^ The position and width of αSMA-expressing or DiI-labeled cells in cultured lungs were measured manually and then compared using ANOVA. For surgical removal and morpholino experiments, corrected total cellular fluorescence (CTCF) was calculated for the αSMA signal surrounding the first branch of the lung. CTCF was calculated by measuring the integrated greyscale density of the αSMA channel and subtracting the product of the area and mean fluorescence of the background. Due to the directional hypothesis of surgical removal and *PITX2*-targetting morpholinos reducing αSMA expression, we used one-tailed t-tests to test for significant differences between experimental and control groups.

## ACKNOWLEDGEMENTS

Princeton High Meadows Environmental Institute to CMN and RM

- NSF (PRFB — DBI-2209090 to AHG; DBI-2305831 to BL)
- NIH (HD111539, HL164861, HD099030, HL166311 to CMN; DE030326 to MR)
- NSERC PGS-D, Canadian Federation of University Women Dr. Margaret McWilliams Predoctoral Fellowship, Princeton University Procter Fellowship, and American Heart Association Predoctoral Fellowship to KG
- American Heart Association Postdoctoral Fellowship to CJP

## AUTHOR CONTRIBUTIONS

AHG, KG, RM, and CMN conceived and designed the study. AHG, KG, MAP, BL, CJP, MR, and WW collected the data. AHG, KG, BJB, JAM, RM, and CMN analyzed the data. AHG, KG, RM, and CMN drafted the manuscript. All authors edited and revised the manuscript.

## Abbreviations

αSMA: α-smooth muscle actin
ATAC-seq: assay for transposase accessible chromatin sequencing
CK: cytokeratins
dpo: days postoviposition
*E*: embryonic day
HH: Hamburger & Hamilton stage
scRNA-seq: single cell RNA sequencing
UMAP: uniform manifold approximation and projection

## Notes

### Competing Interest Statement

The authors have declared no competing interest.

### Summary of Updates

This manuscript was revised to reflect new authors, new data, a new title, and new text.

