## Supplemental Materials for "A vascular origin for pulmonary smooth muscle in the avian lung"

**Supplemental Material**

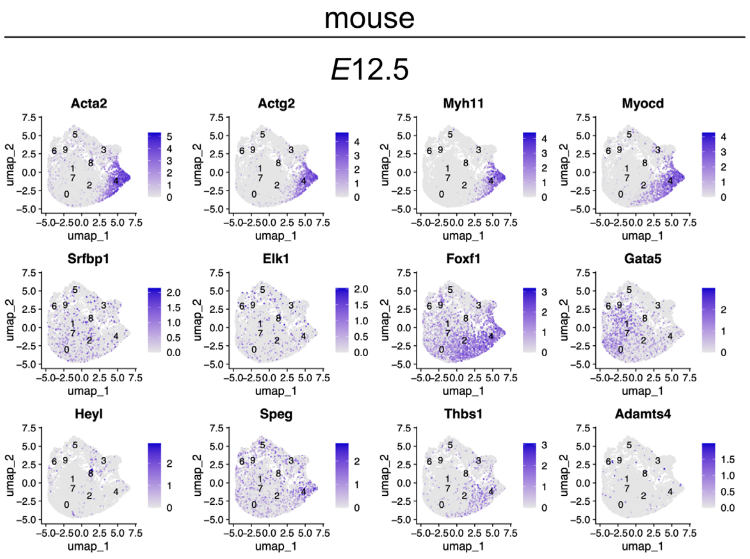


**Supplemental Figure 1. scRNA-seq of smooth-muscle markers in embryonic mouse lung mesenchyme**. UMAPs illustrate cells within mesenchymal clusters of the *E*12.5 mouse lung that express smooth-muscle markers outlined by Jaslove and Nelson (2018).

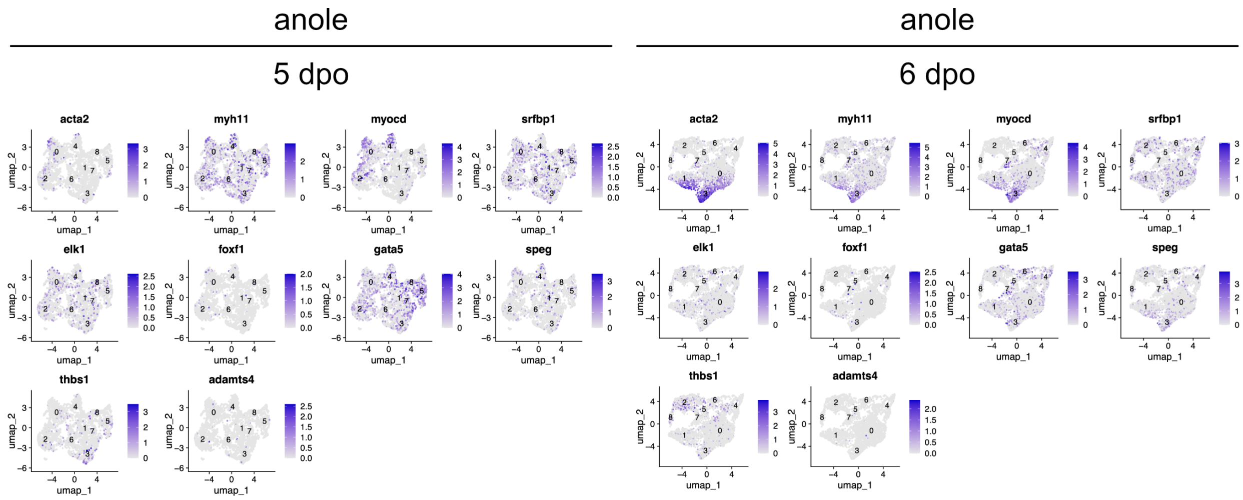


**Supplemental Figure 2. scRNA-seq of smooth-muscle markers in embryonic anole lung mesenchyme**. UMAPs illustrate cells within mesenchymal clusters of 5-dpo and 6-dpo anole lungs that express smooth -muscle markers outlined by Jaslove and Nelson (2018).

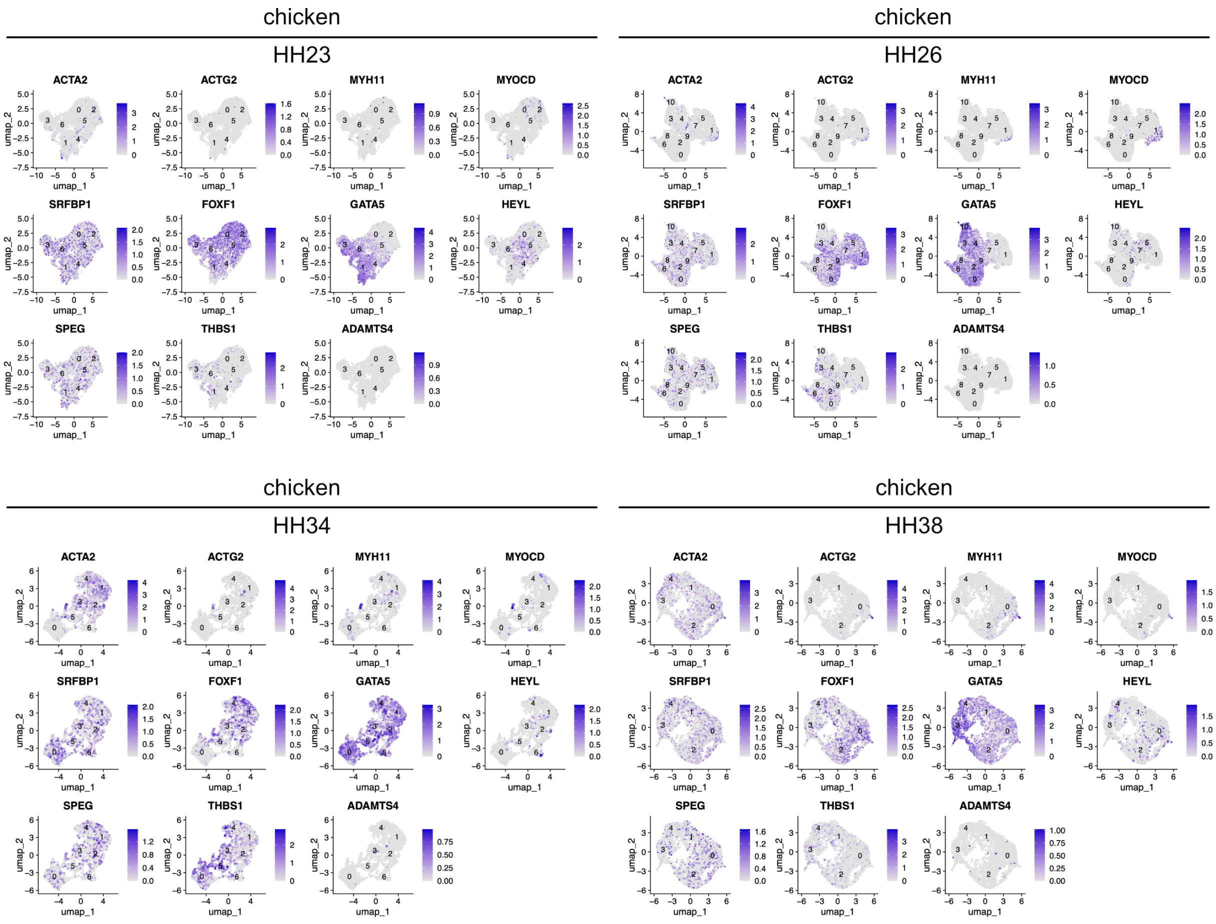


**Supplemental Figure 3. scRNA-seq of smooth-muscle markers in embryonic chicken lung mesenchyme**. UMAPs illustrate cells within mesenchymal clusters of HH23-, HH26-, HH34-, and HH38-stage chicken lungs that express smooth-muscle markers outlined by Jaslove and Nelson (2018).

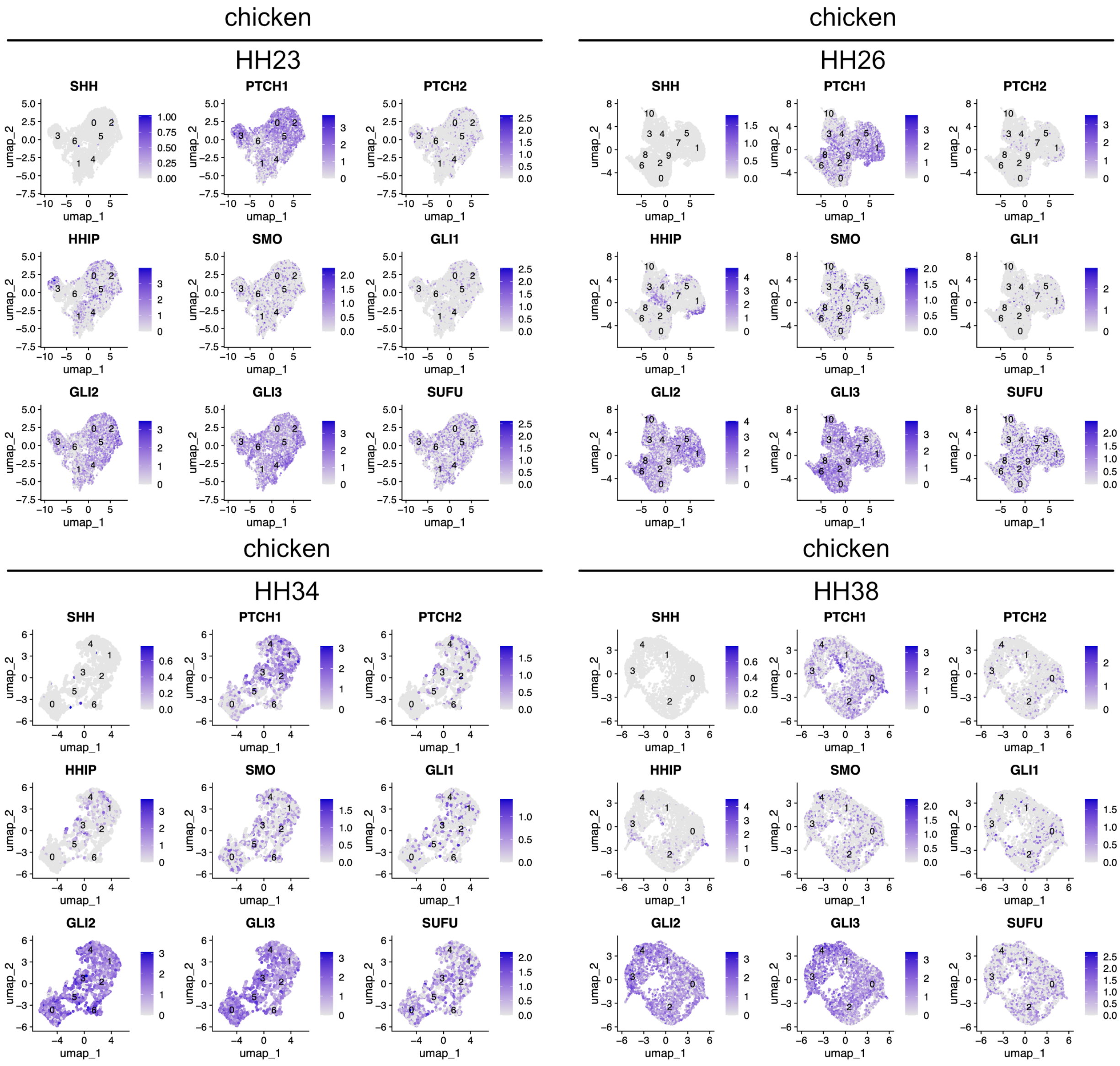


**Supplemental Figure 4. scRNA-seq of sonic hedgehog-pathway genes in embryonic chicken lung mesenchyme**. UMAPs illustrate cells within mesenchymal clusters of HH23-, HH26-, HH34-, and HH38-stage chicken lungs that express SHH-pathway genes.

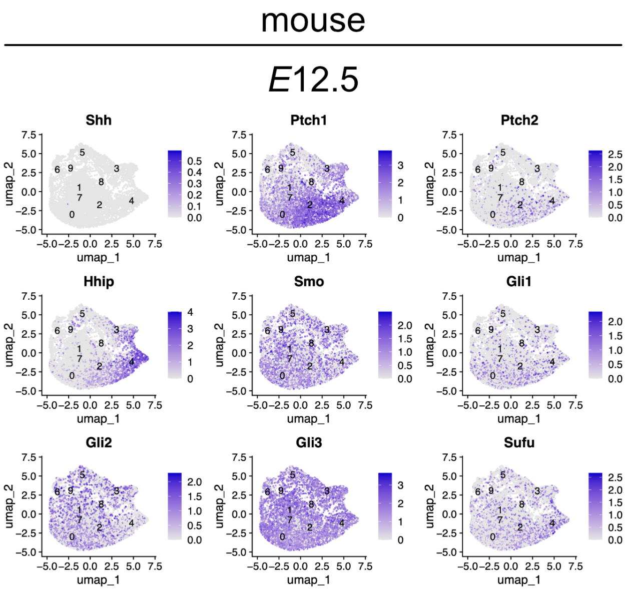


**Supplemental Figure 5. scRNA-seq of sonic hedgehog-pathway genes in embryonic mouse lung mesenchyme**. UMAPs illustrate cells within mesenchymal clusters of the *E*12.5 mouse lung that express SHH-pathway genes.

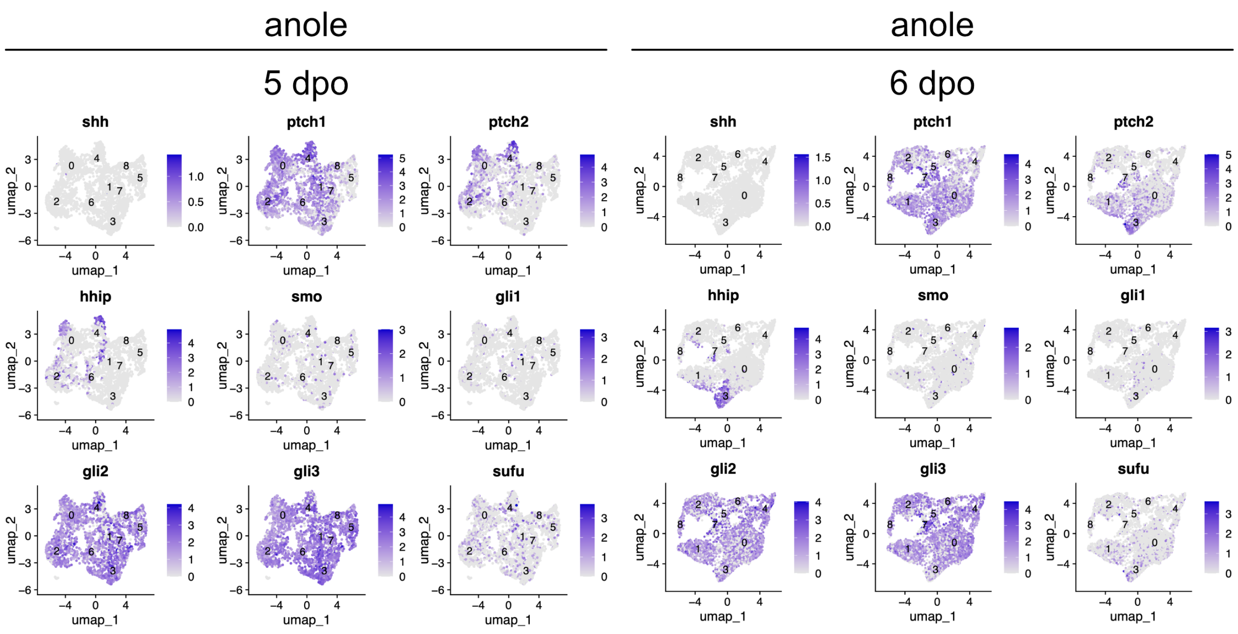


**Supplemental Figure 6. scRNA-seq of sonic hedgehog-pathway genes in embryonic anole lung mesenchyme**. UMAPs illustrate cells within mesenchymal clusters of 5-dpo and 6-dpo anole lungs that express SHH-pathway genes.

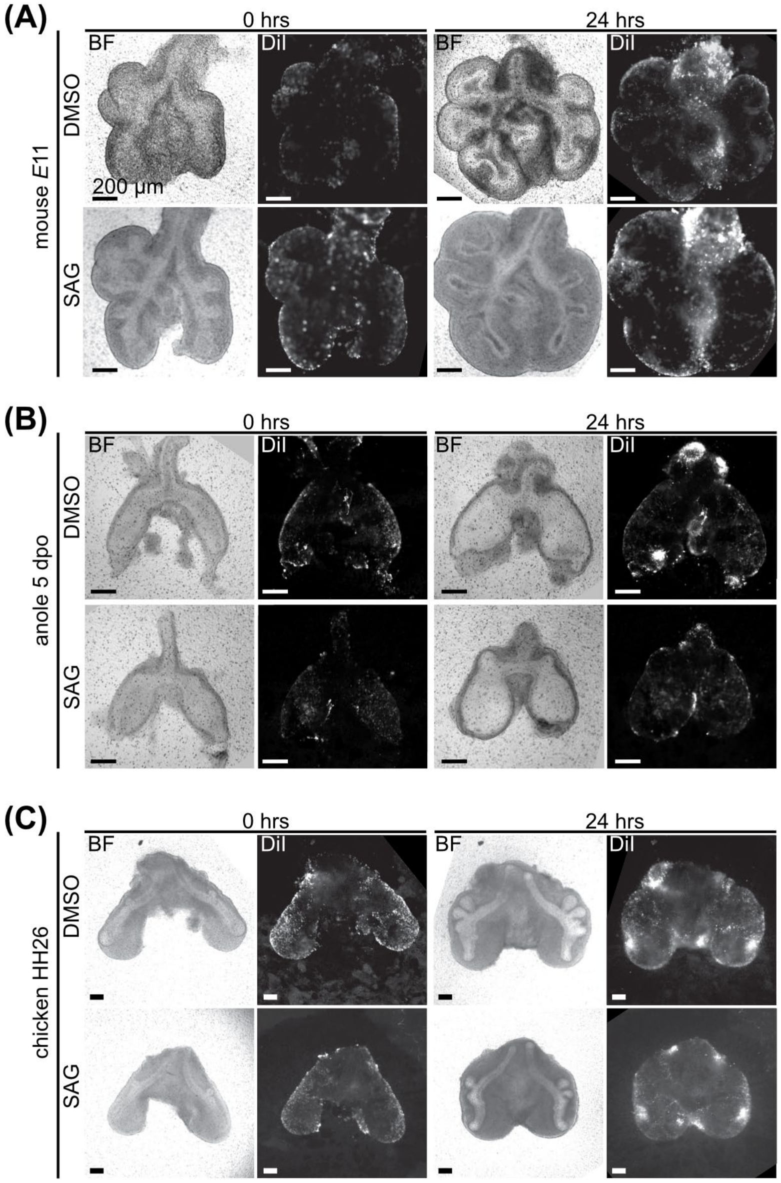


**Supplemental Figure 7. Smoothened agonist and DiI experiments**. Brightfield (BF) and fluorescence images of mouse lungs isolated at *E*11 (**A**), anole lungs isolated at 5-dpo (**B**), or chicken lungs isolated at HH26 (**C**), incubated with DiI to label the mesothelial surface, and then cultured *ex vivo* for 24 hours in the presence of DMSO or SAG. Scale bars = 200 μm.

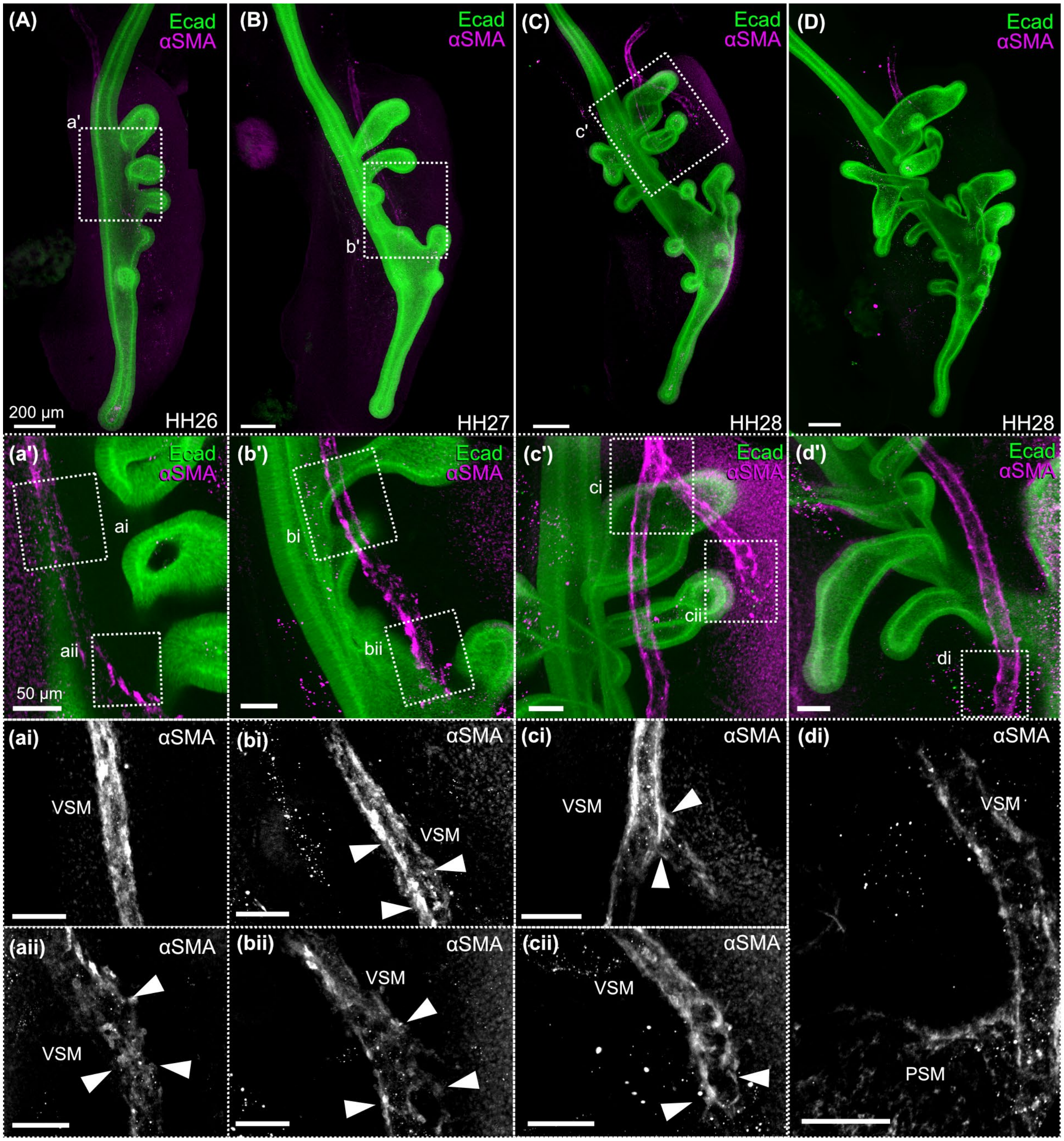


**Supplemental Figure 8. Immunofluorescence analysis of smooth muscle development in duck lungs**. Images show immunofluorescence for alpha smooth muscle actin (αSMA, magenta) and E-cadherin (Ecad, green) in developing lungs of Pekin duck (*Anas platyrhynchos*) at HH26 (**A**), HH27 (**B**), and HH28 (**C**, **D**). Scale bars = 200 µm. Inset images focus on vascular smooth muscle (VSM). Inset image in panel **D** includes pulmonary smooth muscle (PSM) surrounding airways adjacent to VSM. White arrows point to irregular organization of αSMA+ cells, similar to that of chicken VSM in Figure 4G. Scale bars = 50 µm.

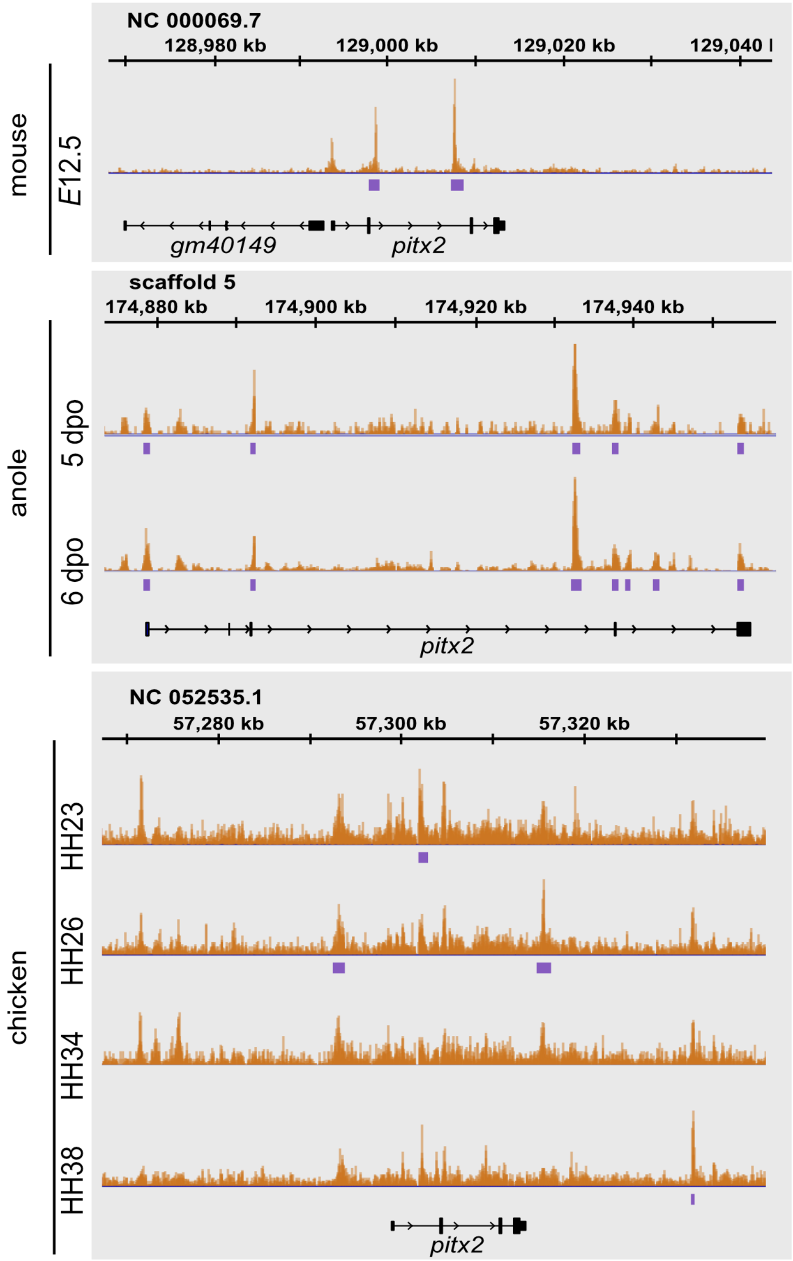


**Supplemental Figure 9**. **ATAC-seq of the *PITX2* regulatory locus in mouse, anole, and chicken lungs.** Pictured are open chromatin pileups (orange) and narrow peaks (purple) surrounding *PITX2* regulatory regions.

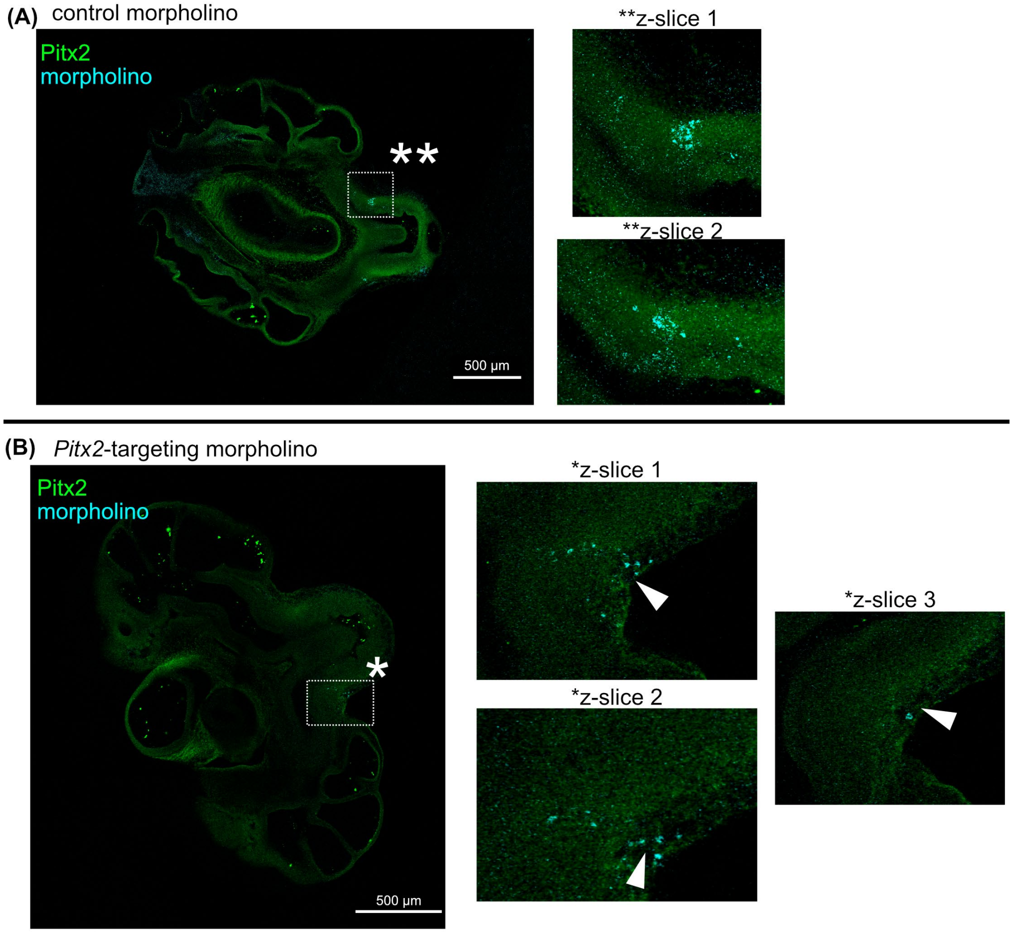


**Supplemental Figure 10. *PITX2*-targeting morpholino experiments**. Immunofluorescence for PITX2 (green) and fluorescein-conjugated morpholinos (cyan) in HH26 + 48 hour *ex vivo* culture chicken lungs. In the control morpholino replicates (**A**), PITX2 signal is not attenuated surrounding morpholinos. In *PITX2*-targeting morpholino replicates (**B**), PITX2 signal is absent surrounding morpholinos (white arrows). Scale bars = 500 µm.
